# Conservation and Divergence in the Asexual Sporulation Gene Regulatory Network Across a Genus of Filamentous Fungi

**DOI:** 10.1101/331868

**Authors:** Ming-Yueh Wu, Matthew E. Mead, Mi-Kyung Lee, Sun-Chang Kim, Antonis Rokas, Jae-Hyuk Yu

## Abstract

Asexual sporulation is fundamental to the ecology and lifestyle of filamentous fungi and can facilitate both plant and human infection. In *Aspergillus*, the production of asexual spores is primarily governed by the BrlA→AbaA→WetA regulatory cascade. The final step in this cascade is controlled by the WetA protein and not only governs the morphological differentiation of spores but also the production and deposition of diverse metabolites into spores. While WetA is conserved across the genus *Aspergillus*, the structure and degree of conservation of the *wetA* gene regulatory network (GRN) remains largely unknown. We carried out comparative transcriptome analyses between *wetA* null mutant and wild type asexual spores in three representative species spanning the diversity of the genus *Aspergillus*: *A. nidulans, A. flavus*, and *A. fumigatus*. We discovered that WetA regulates asexual sporulation in all three species via a negative feedback loop that represses BrlA, the cascade’s first step. Furthermore, ChIP-seq experiments in *A. nidulans* asexual spores suggest that WetA is a DNA-binding protein that interacts with a novel regulatory motif. Several global regulators known to bridge spore production and the production of secondary metabolites show species-specific regulatory patterns in our data. These results suggest that the BrlA→AbaA→WetA cascade’s regulatory role in cellular and chemical asexual spore development is functionally conserved, but that the *wetA*-associated GRN has diverged during *Aspergillus* evolution.

## Importance

The formation of resilient spores is a key factor contributing to the survival and fitness of many microorganisms, including fungi. In the fungal genus *Aspergillus*, spore formation is controlled by a complex gene regulatory network that also impacts a variety of other processes, including secondary metabolism. To gain mechanistic insights into how fungal spore formation is controlled across *Aspergillus*, we dissected the gene regulatory network downstream of a major regulator of spore maturation (WetA) in three species that span the diversity of the genus: the genetic model *A. nidulans*, the human pathogen *A. fumigatus*, and the aflatoxin producer *A. flavus*. Our data shows that WetA regulates asexual sporulation in all three species via a negative feedback loop and likely binds a novel regulatory element we term the WetA Response Element (WRE). These results shed light on how gene regulatory networks in microorganisms control important biological processes and evolve across diverse species.

## Introduction

The ability to produce numerous asexual spores is one of the key factors contributing to the fecundity and fitness of filamentous fungi. Fungal asexual spores are highly efficient for genome protection, survival, and propagation. Spores are also the primary means of infecting host organisms for many pathogenic fungi (1). Importantly, in some filamentous fungi, morphological development is coordinated with the production of secondary metabolites with toxic and antibiotic properties (2–4).

Asexual development (conidiation) in the fungal class Eurotiomycetes results in the formation of mitotically derived asexual spores known as conidiospores or conidia. As asexual sporulation is widespread among fungi, it represents a simple, highly tractable system for understanding how gene regulatory networks (GRNs) evolve in microbial eukaryotes and how this evolution has influenced developmental and metabolic phenotypes.

Members of the genus *Aspergillus* are ubiquitous in most environments, and include various beneficial, pathogenic, and/or toxigenic species (5) All aspergilli produce conidia as the main means of dispersion and infection. Importantly, the asexual development and the production of certain secondary metabolites including mycotoxins are intimately associated (2).

The three distantly related species *Aspergillus nidulans, Aspergillus flavus*, and *Aspergillus fumigatus*, whose pairwise levels of genome similarity are similar to the genomic similarity between the human and fish genomes (6), form distinct conidiophores with varying sizes of conidia. The regulatory mechanisms of conidiation have been extensively studied in *A. nidulans* (7–23). The regulatory genes can be divided into central regulators, upstream activators, negative regulators, light-dependent regulators, and the *velvet* regulators (24, 25). The central genetic regulatory cascade BrlA→AbaA→WetA is present in *Aspergillus* and governs both conidiation-specific GRNs and the resulting morphological pathway of conidiation (Fig 1A) (22, 24, 26). BrlA is a C_2_H_2_-zinc-finger type transcription factor (TF), which recognizes and interacts with BrlA Response Elements (BRE) (Fig 1B) (27, 28). The *brlA* gene is expressed in the early phase of conidiation and mediates vesicle formation and budding-like cell growth (11). The *abaA* gene is activated by BrlA and regulates the formation of metulae and phialides. Similar to BrlA, AbaA is a TF, containing an TEA/ATTS DNA binding motif and a potential leucine zipper that recognizes the AbaA Response Elements (AREs) (Fig 1B) (29). The *wetA* (wet-white A) gene, activated by AbaA, functions in the late phase of conidiation that completes sporogenesis. The BrlA→AbaA→WetA central regulatory cascade acts in concert with other genes to control conidiation-specific gene expression and determine the order of gene activation during the cellular and chemical development of spores. The WetA protein plays a pivotal role in the coordinated control of *Aspergillus* conidiogenesis; however, the precise molecular mechanisms of WetA function have been unknown. WetA is highly and broadly conserved in *Ascomycetes* (8, 10–13, 15–23, 26, 30), plays an essential role in the synthesis of crucial conidial wall components, and makes the conidia both impermeable and mature (20, 21, 30). The *Aspergillus* WetA proteins have a conserved ESC1/WetA-related domain (PTHR22934: SF23) with putative DNA-binding functions, a predicted transcription activation domain (TAD) (31), and a nuclear localization signal (NLS) (32, 33) near the C terminus (16, 34), suggesting that WetA is likely a DNA-binding TF (30) (Fig 1B). As summarized in Table 1, the deletion of *wetA* results in a plethora of conidial defects, including the formation of colorless conidia that undergo autolysis in *A. nidulans* (10–13, 20–22, 26), *A. fumigatus* (8, 15), *A. oryzae* (18), and *A. flavus* (30). The metabolism and expression of several conidial components are perturbed in the Δ*wetA* conidia, leading to reduced stress tolerance and spore viability (8, 30).

**Table 1.**
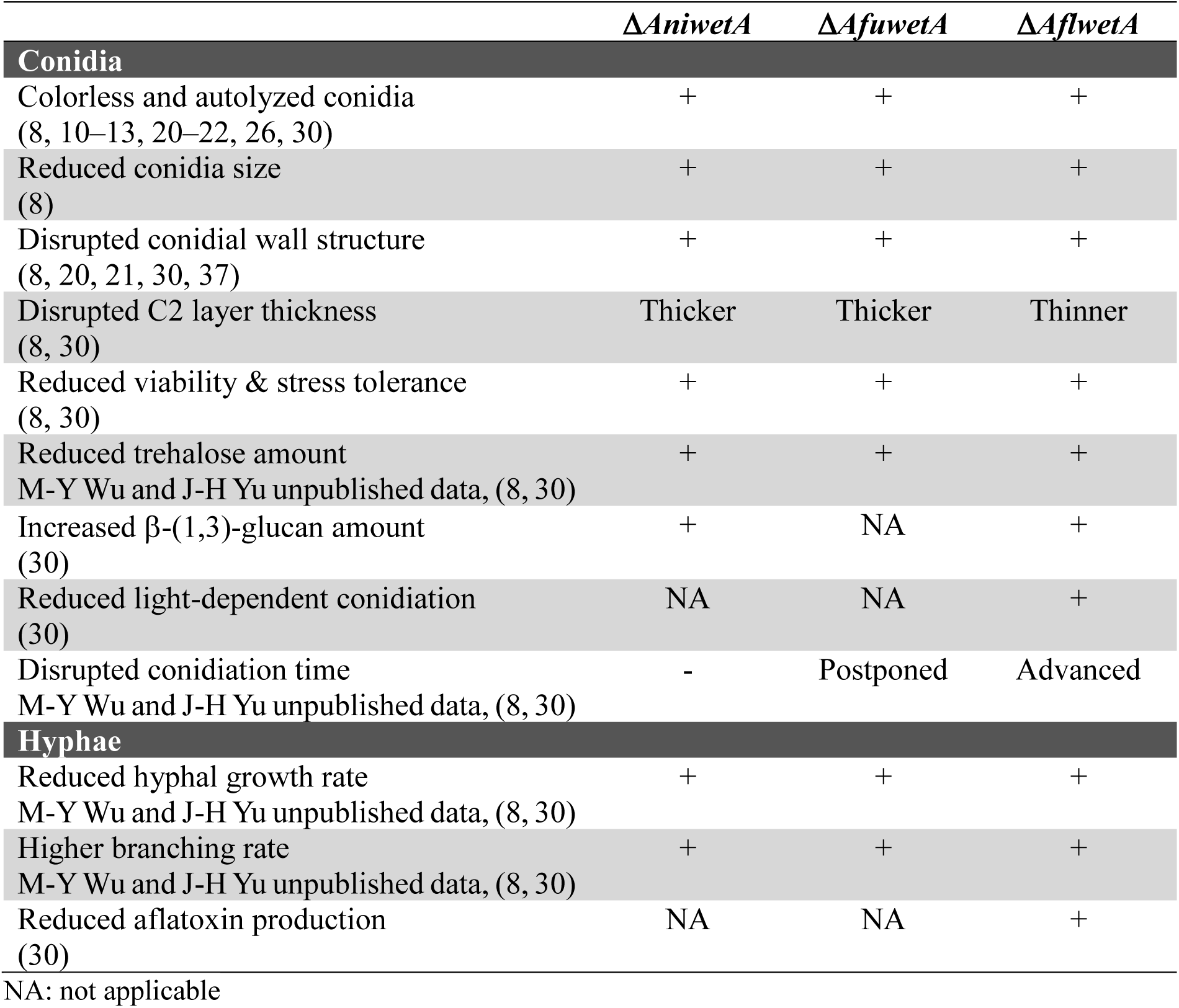
The roles of WetA in three *Aspergillus* species

**Fig 1.**
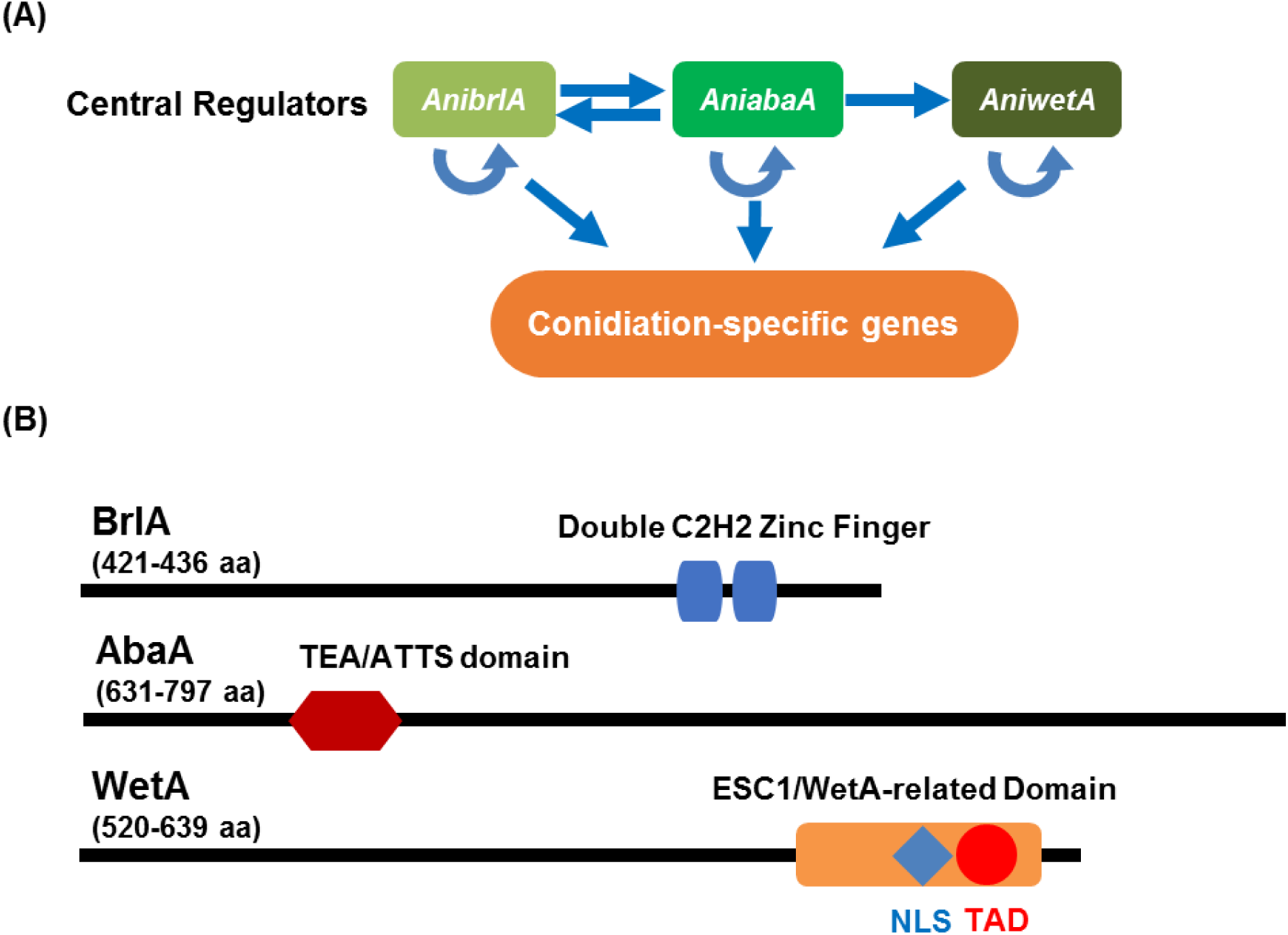
The central regulatory pathway of *Aspergillus* conidiation. (A) A cartoon depiction of genetic interactions of the central regulators in *A. nidulans* conidiogenesis. The central regulators cooperatively activate the conidiation-specific genes responsible for the morphogenesis of conidiophores. (B) The predicted protein architectures for the three conserved central regulators of conidiation in *A. nidulans, A. fumigatus*, and *A. flavus*. The blue box and the red hexagon represents the C_2_H_2_ zinc finger domain and TEA/ATTS domain in BrlA and AbaA, respectively, and were identified in a blastP (version 2.6.0) search (35). The red circle represents a putative transcription activation domain (TAD), which was predicted by 9aaTAD using the “Less stringent Pattern” setting (31). The blue diamond represents the nuclear localization signal (NLS) predicted by NLStradamus using the 4 state HMM static model (32). The orange rectangle represents the ESC1/WetA-related domain (PTHR22934) predicted by the PANTHER classification system (36).

In this report, we have investigated the structure and degree of conservation of the BrlA→AbaA→WetA central regulatory cascade of *Aspergillus* conidiation and the broader *wetA* GRNs in three representative *Aspergillus* species: the genetic model *A. nidulans*, the mycotoxin producer *A. flavus*, and the human pathogen *A. fumigatus*. Specifically, we carried out comparative transcriptome analyses between *wetA* null mutant and wild type (WT) asexual spores in the three species. We also investigated the WetA-chromatin interaction in asexual spores via ChIP-seq in *A. nidulans* spores, which enabled us to identify the consensus WetA-DNA binding sequence. Further comparative genome-wide analyses revealed that the WetA-associated GRN has diverged during *Aspergillus* evolution, uncovering important yet unexplored regulatory networks of asexual sporulation and metabolic remodeling in *Aspergillus*. Our findings provide the first clear and systematic dissection of the evolutionarily conserved WetA developmental regulator governing the diverged processes of cellular differentiation, chemical development, and cell survival across a genus of filamentous fungi.

## Results

### Conserved and diverged roles for WetA in the control of gene expression in *Aspergilli*

To investigate the conserved and divergent regulatory roles WetA plays in the three *Aspergillus* species, we carried out comprehensive analyses of gene expression differences between the WT and *wetA* null mutant conidia. We found that WetA plays a broad regulatory role in conidia in all three *Aspergillus* species; approximately 52%, 57%, and 43% of all genes showed differential accumulation of mRNAs in the Δ*wetA* conidia in comparison to WT conidia in *A. nidulans, A. fumigatus*, and *A. flavus*, respectively (Table 2). Among the differentially expressed genes (DEGs), 46%, 48%, and 50% were underexpressed and 54%, 52%, and 50% were overexpressed in the Δ*wetA* conidia compared to the WT conidia in *A. nidulans, A. fumigatus*, and *A. flavus*, respectively (Table 2).

**Table 2.** Summary of DEGs in the three *Aspergillus* Δ*wetA* conidia

|  | *A. nidulans* | *A. fumigatus* | *A. flavus* |
| --- | --- | --- | --- |
| Unaffected genes | 5,246 (48%) | 4,374 (43%) | 7,730 (57%) |
| **DEG** | **5,742 (52%)** | **5,756 (57%)** | **5,755 (43%)** |
| Overexpressed in $\Delta wetA$ | 3,107 (28%) | 2,996 (30%) | 2,899 (21%) |
| Underexpressed in $\Delta wetA$ | 2,635 (24%) | 2,758 (27%) | 2,856 (21%) |
| **Total** | **10,988** | **10,130** | **13,485** |

Functional category analysis was carried out by determining Gene Ontology (GO) terms that were enriched in DEGs. Specifically, the biological process GO categories that were enriched in the Δ*wetA* conidia included “asexual sporulation”, “secondary metabolic process”, and “toxin biosynthetic process”. Moreover, over 70% of all genes in the cellular component GO category, “fungal-type cell wall”, were also regulated in each species. These top enriched GO categories are consistent with the phenotypes of the Δ*wetA* mutants, suggesting that WetA plays key a role in carbohydrate metabolism, secondary metabolism, and conidial wall integrity (30). To explore the conserved and diverged regulatory roles of WetA, we examined the mRNA expression profiles of orthologous groups of genes (orthogroups) in the three *Aspergillus* genomes. In total, 8,978 orthogroups were identified and 6,466 of these contained orthologs in all three species. Of the 8,978 total orthogroups, 7,301 (81%) had at least one gene that showed differential expression in the Δ*wetA* conidia, but only 1,294 orthogroups show consistent WetA-regulation (i.e., all orthologs in the group were either overexpressed or underexpressed). The enriched GO categories of the 1,294 orthogroups whose genes showed the same differential expression pattern suggest that WetA is functionally conserved in controlling stress response, pigmentation, spore trehalose formation, cell wall organization, and cellular development, which are also consistent with the phenotypes observed in Δ*wetA* strains. In contrast, the remaining 6,007 WetA-regulated orthogroups show divergent differential expression patterns, implying that a substantial portion of the WetA-controlled GRNs has functionally diverged among the three species. Furthermore, of the 6,466 *Aspergillus* orthogroups that contain orthologs in all three species, only 788 exhibited a conserved pattern of differential expression (i.e., all genes were either overexpressed or underexpressed in the Δ*wetA* conidia in all three species) (Fig 2, Table S2).

**Fig 2.**
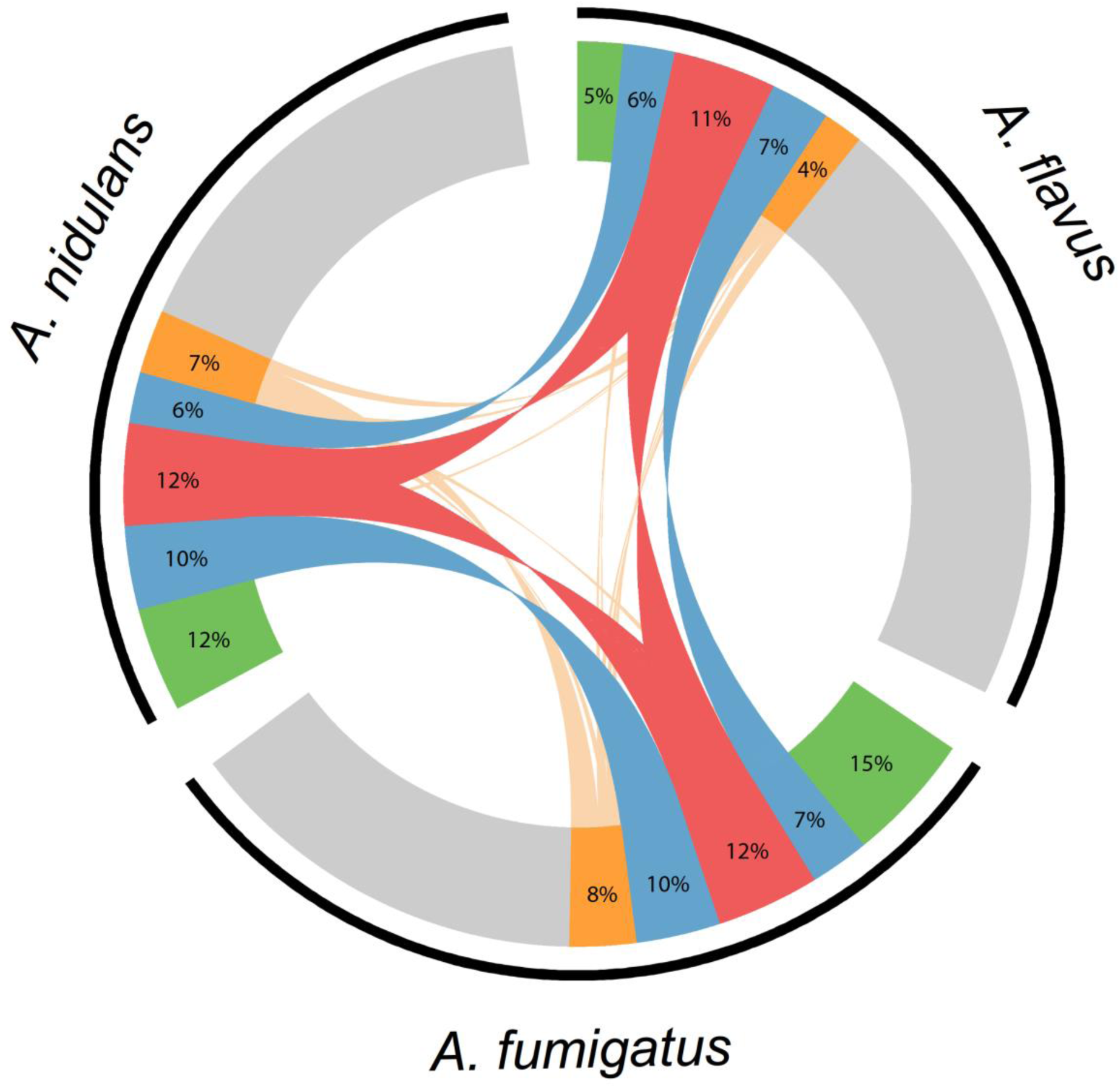
Overview of the WetA-regulated-orthologs in *A. nidulans, A. fumigatus*, and *A. flavus*. The 6,466 genes belonging to an orthogroup that possessed at least one member from *A. nidulans, A. fumigatus*, and *A. flavus* are represented by the black arcs next to their respective species labels. Gray, orthologs whose expression did not change between Δ*wetA* and WT conidia. Green, orthologs that were differentially expressed in only one species. Blue, genes that showed the same differential expression pattern in two out of the three species. Red, genes that showed the same differential expression pattern in all three species. Orange, genes that showed a divergent differential pattern in two or more species. Lines connect expressed genes from the same orthogroup.

## WetA-regulated genes involved in asexual development, signal transduction, and conidial integrity are divergently regulated among *Aspergilli*

To explore the conserved and diverged molecular roles of WetA in conidiation in the three species, we examined mRNA levels of genes related to asexual development, signal transduction, and conidial integrity (Fig 3, Table S3), phenotypes previously implicated to be controlled by WetA (8, 10–13, 20–22, 26, 30, 37).

**Fig 3.**
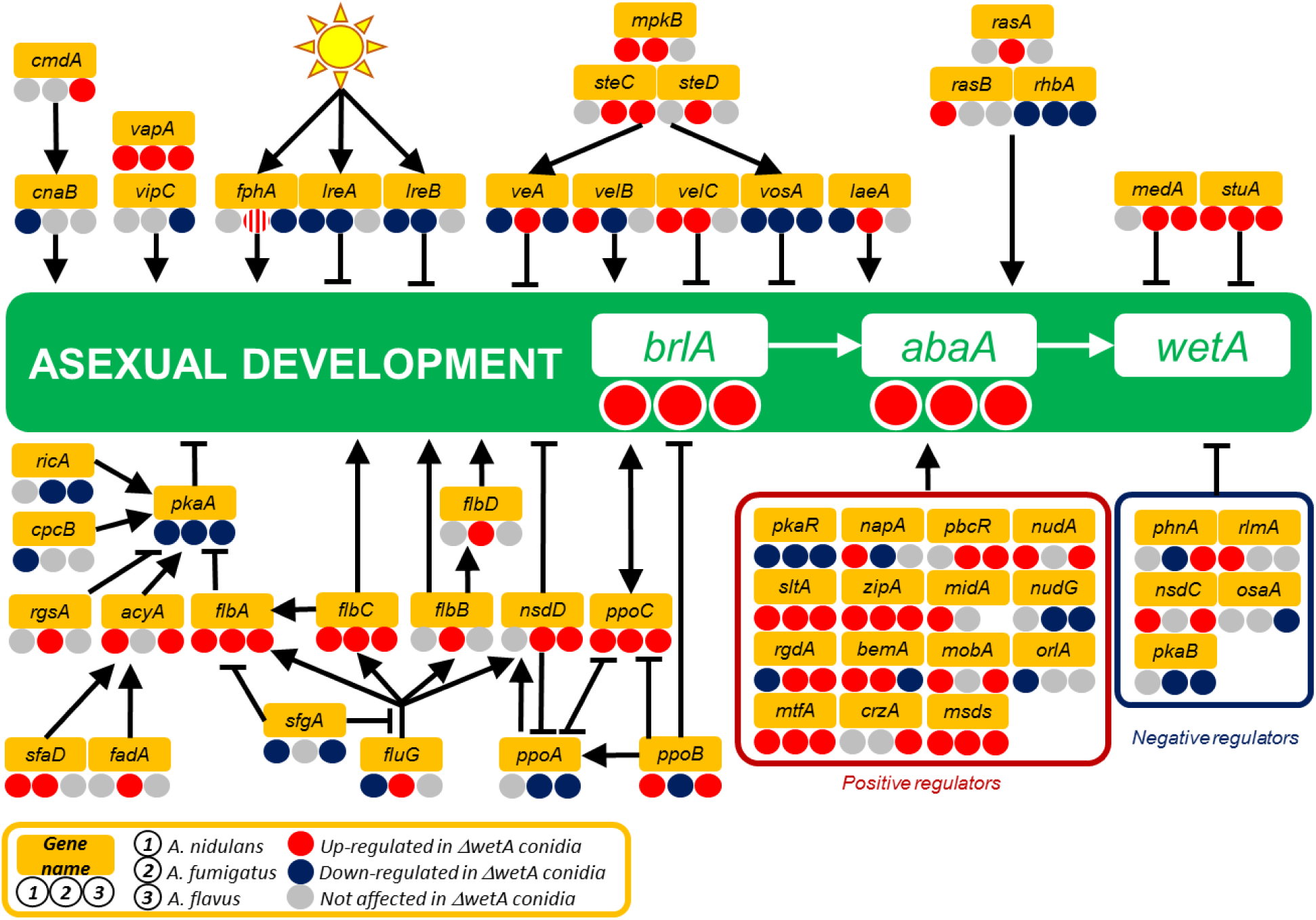
WetA-mediated regulation of asexual development in the three Aspergillus species. Schematic diagram of the WetA-mediated regulatory model of conidiation. Genes with increased, decreased, and unaffected mRNA levels in the Δ*wetA* conidia are labeled with red (WetA-inhibited), blue (WetA-activated), and grey (not affected by WetA) circles and the WetA-regulatory effects in the Δ*AniwetA*, Δ*AfuwetA*, and Δ*AflwetA* conidia are listed under the gene name from left to right, respectively. There are two orthologs of *fphA* in *A. fumigatus*, one of which is WetA-inhibited and the other is not regulated by WetA.

Our data show that WetA negatively regulates asexual development in conidia produced by species across the genus *Aspergillus* via a negative feedback loop that represses the pathway’s upstream regulator, *brlA*. Specifically, both *brlA* and *abaA* expression are increased in the Δ*wetA* conidia relative to WT in all three species (Figure 4). However, to achieve the conserved repression of *brlA* and *abaA* mRNA accumulation, WetA shows species-specific regulatory effects on *brlA* upstream regulatory networks. For example, in the *velvet* protein family and complex, *vosA* was consistently underexpressed in the three Δ*wetA* conidia, but the WetA effects on *veA, velB, velC*, and *laeA* expression are not conserved in each species. Similarly, the light-dependent regulators were differentially regulated by WetA. The blue-light-dependent regulators *lreA* and *lreB* were unaffected in the Δ*AflwetA* conidia but repressed in both the Δ*AniwetA* and Δ*AfuwetA* conidia. Taken together, the WetA-mediated feedback repression of asexual development is functionally conserved across the genus *Aspergillus* but the specific GRNs appear to have diverged during the evolution of the genus *Aspergillus*.

Our previous study showed that *Afl*WetA is involved in regulating G-protein regulatory pathways (30). Expanding this analysis in the three *Aspergillus* species showed that *gprC, gprF,gprG, nopA, flbA*, and *pkaA* were consistently differentially regulated in the Δ*wetA* conidia, while other members in the G-protein regulatory pathways were either not affected by WetA or showed species-specific regulatory patterns in the Δ*wetA* conidia (Table S4).

WetA is involved in other signal transduction pathways. Total 110, 126, and 92 kinase-encoded genes were differentially expressed in the Δ*AniwetA*, Δ*AfuwetA*, and Δ*AflwetA* conidia respectively; however, only 21 of them were consistently over-or underexpressed in the Δ*wetA* conidia of all three species (Table S5). Similarly, 132, 153, and 142 putative TF-encoded genes in each species were differentially expressed in Δ*AniwetA*, Δ*AfuwetA*, and Δ*AflwetA* conidia respectively; however, only 32 were consistently over- or underexpressed in all three of the Δ*wetA* conidia (Table S5).

We further investigated the mRNA levels of the genes in the secondary metabolite gene (SMG) clusters in each species (30, 38, 39) (Table S7). In total, 96% (64/67), 100% (33/33), and 92% (68/74) of SMG clusters in the Δ*AniwetA*, Δ*AfuwetA*, and Δ*AflwetA* conidia, respectively, had at least one gene that showed altered mRNA expression levels (Table 3). One of the SMG backbone genes, *wA*, is conserved in all three species and it encodes a polyketide synthase (PKS) necessary for the formation of a key conidial pigment (40). Previous studies showed that *wA* is activated by WetA (20), consistent with the colorless conidia phenotype of the Δ*wetA* mutants. Although *wA* was underexpressed in the Δ*AniwetA* and Δ*AflwetA* conidia as expected, it was overexpressed in Δ*AfuwetA*, suggesting that the regulation of the conidial pigmentation pathway in *A. fumigatus* differs from that in the other two species.

**Table 3.**
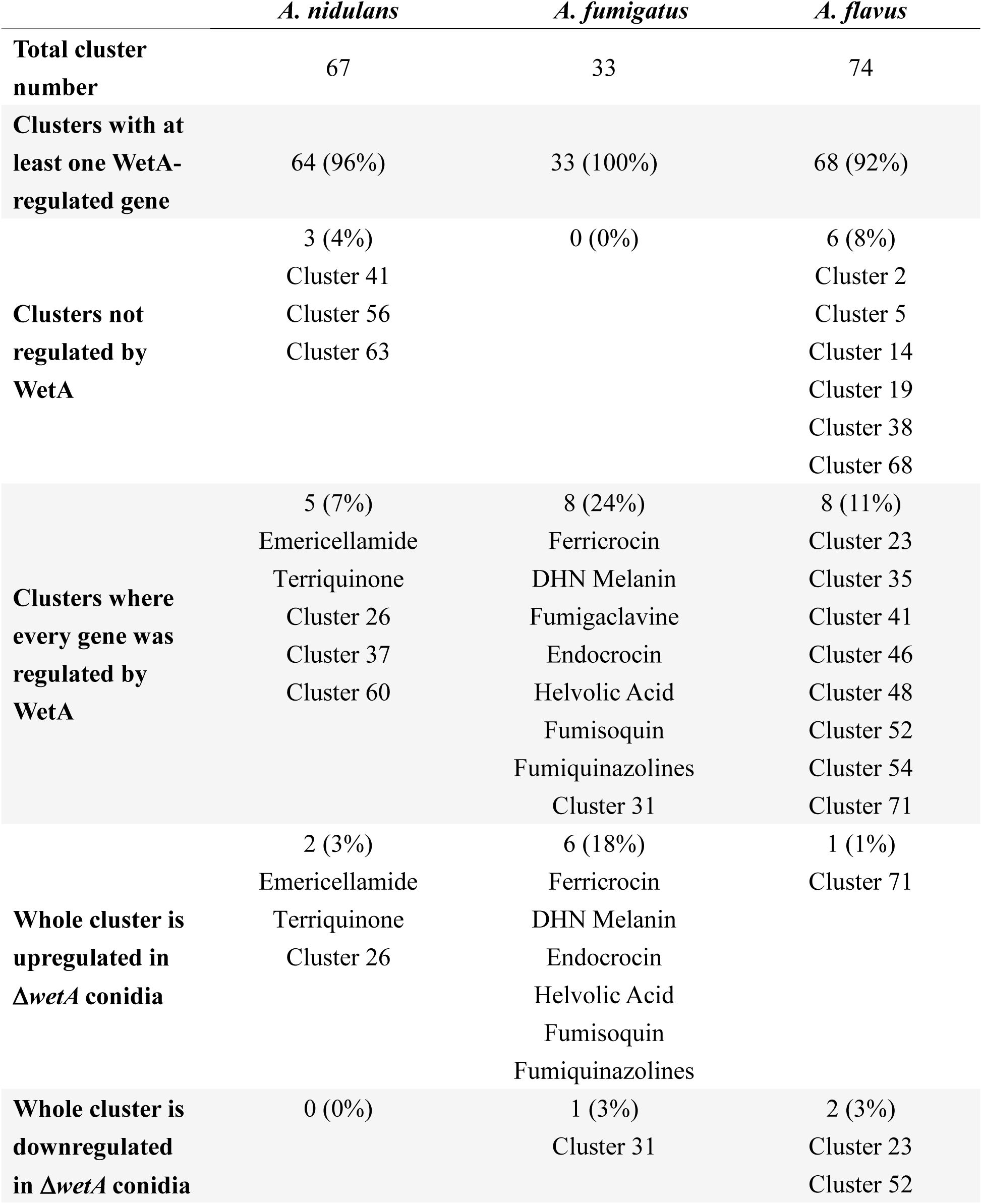
WetA-mediated SMG regulation

Finally, we examined the expression levels of genes involved in conidia content and conidial wall integrity. Most of the DEGs associated with trehalose biosynthesis were underexpressed in the Δ*wetA* conidia in all three species, while *treA*, involved in trehalose degradation, was overexpressed in the Δ*AniwetA* and Δ*AfuwetA* conidia but underexpressed in the Δ*AflwetA* conidia (Fig 4). Loss of *wetA* resulted in overexpression of almost all genes involved in the biosynthesis of chitin and β-(1,3)-glucan, but genes involved in the biosynthesis and degradation of α-(1,3)-glucan were both overexpressed and underexpressed relative to WT. (Fig 4).

**Fig 4.**
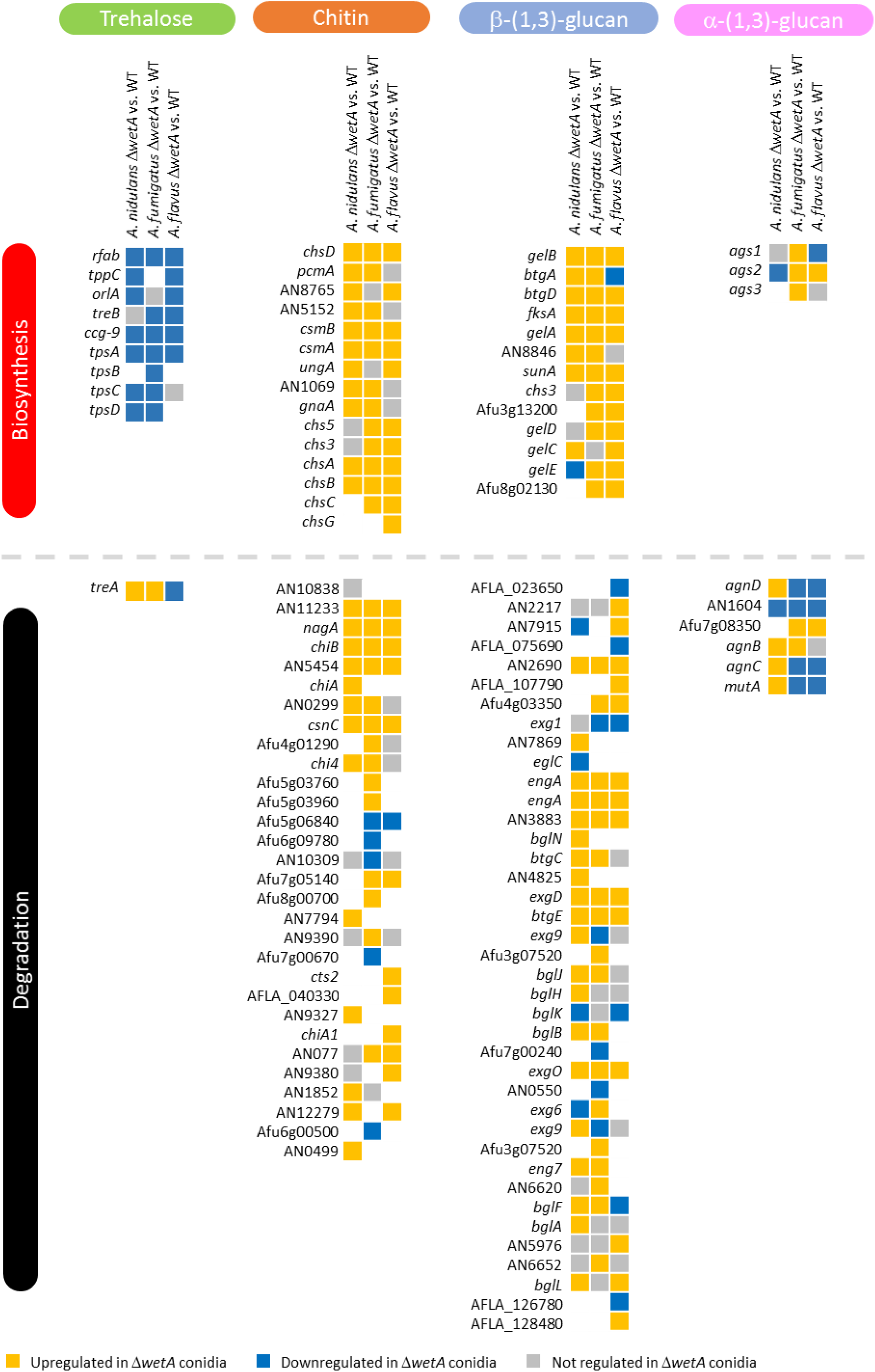
WetA-regulatory effects on trehalose, chitin, β-(1,3)-glucan, and α-(1,3)-glucan metabolism in *Aspergillus* species.

Moreover, our results show that WetA is a key regulator of hydrophobins, DHN-melanin biosynthesis, and pyomelanin biosynthesis. Somewhat unexpectedly, although we observed the conserved “wet” and “white” phenotypes of the Δ*wetA* conidia in all three species, all of the genes proposed to be related to the “wet” (hydrophobin) and “white” (DHN-melanin and pyomelanin) phenotypes were overexpressed in the Δ*AfuwetA* conidia (Fig 5).

**Fig 5.**
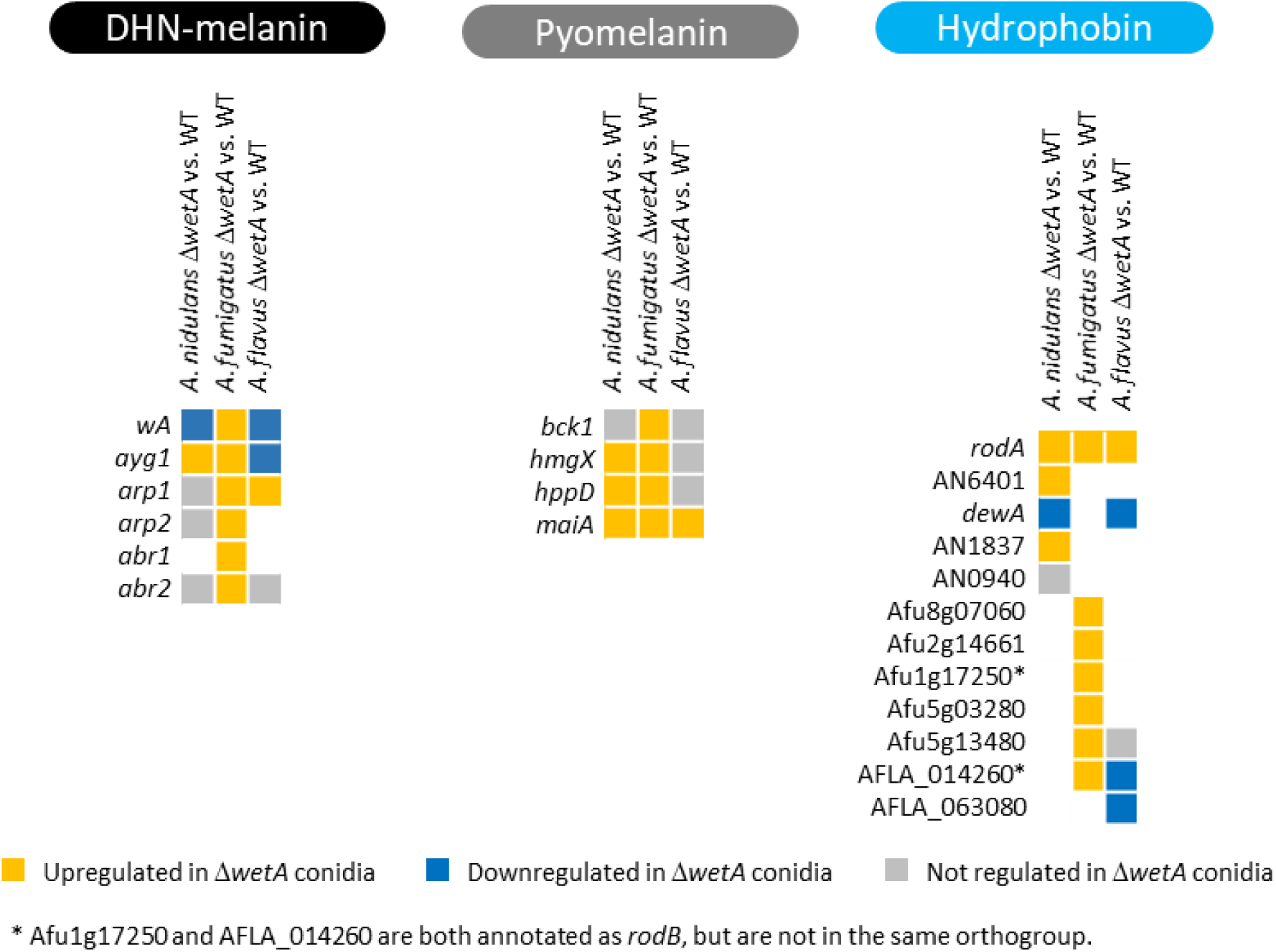
WetA-regulatory effects on DHN-melanin, pyomelanin, and hydrophobin biosynthesis in *Aspergillus* species. In summary, our RNA-seq analyses suggest that WetA exerts broad regulatory effects in conidiation by controlling about half of the transcriptome in each of the three *Aspergillus* species we tested. Even though WetA-mediated regulation results in similar phenotypes and pathways being regulated across the three species, the nature of that regulation is different when comparing individually regulated genes among the species. These results suggest that, although the WetA-mediated GRNs have diverged during the evolution of *Aspergillus*, their regulatory logic appears to have remained conserved.

### Identification of WetA response elements (WREs)

To better understand WetA regulatory mechanisms in conidia, we carried out chromatin immunoprecipitation (ChIP) experiments, followed by high-throughput sequencing of the enriched DNA fragments (ChIP-seq) in the *A. nidulans* conidia. We identified 157 peaks from two independent ChIP-seq experiments, using a False Discovery Rate (FDR)?cutoff of less than or equal to?0.001 and a Fold Change (FC, sample tag counts divided by input tag counts) cutoff greater than or equal to 2. Of the 157 peaks, 135 were located in at least one of the following: a protein coding region, an intron, an upstream region, or a downstream region (Table S8). Upstream and downstream regions were defined as locations within 1.5 kb of the translation start or stop site, respectively. Many peaks were located within multiple features due to the condensed nature of the *A. nidulans* genome; therefore, 212 genes were considered “peak-associated”. Only a few peaks were located within protein coding regions (18) or introns (5); however, 105 peaks were in upstream regions and 59 peaks were in downstream regions. Of the 212 peak-associated genes, 139 showed differential expression in the *A. nidulans* RNA-seq dataset. Multiple previously described genes are in the list of peak-associated genes, including *flbA, mtfA, nopA, velB, sfaD, wetA, vosA, hsp70, srbA*, and *tpsA* (Table S8).

A putative WRE was predicted by MEME-ChIP (41). The 100 bp surrounding the summits of all peaks was used as input for the MEME-ChIP analysis. The only statistically significant motif identified (E-value = 8.8e-8) was 5’-CCGYTTGCGGC-3’ and it exists in the upstream region of *AniwetA*. Potential *Ani*WetA-recognized regions were identified by searching for the predicted motif in the upstream regions of ORFs in the *A. nidulans* genome with FIMO (42). In total, 2,217 genes were predicted to contain the WRE within their upstream regions in *A. nidulans* (Table S9). Functional analysis shows that many biological processes were enriched in these potential *Ani*WetA targeted genes, including “trehalose metabolic process” and “cell wall mannoprotein biosynthetic process”, consistent with what is known about WetA function in conidiation.

To investigate the expression profile of potential *Ani*WetA target genes in conidia, data from the transcriptomic analysis was utilized. In total, 1,176 WRE-containing genes were differentially expressed in *A. nidulans*, including 2 G-protein signaling pathway-associated genes, 22 conidial integrity-associated genes, 14 putative kinase-encoding genes, 22 putative transcription factor-encoding genes, 5 SMG backbone genes, and 11 asexual development-associated genes (Table 4).

**Table 4.** WetA targeted DEGs in Δ*AniwetA* conidia

|  |  |
| --- | --- |
| **G-protein pathway** | *gprC, nopA* |
| **Conidial integrity** | *tppC, ccg-9, tpsC, treA, chsA, chiA, btgD, eglC, engA, btgC, eng7, bglA, ags2, agnD, agnC, hmgX, hppD, maiA, dewA, AN0499, AN1069, AN1837* |
| **Kinase** | *rio2, aromA, nimO, isr1, teaR, pho80, ffkA, panK, nimX, AN3619, AN8213, AN8843, AN10188, AN10551* |
| **TF** | *fcr1, aflR, dbaA, cpcA, vosA, mdpE, wetA, zapA, AN6295, AN1217, AN0817, AN3502, AN3769, AN0094, AN4773, AN6790, AN8111, AN8355, AN8949, AN11169, AN0388, AN10550* |
| **SMG backbones** | *apdA, inpB, AN0016, AN1242, AN9129* |
| **Asexual development** | *ams1, chsB, cnaB, cpcB, dewA, gprC, odeA, tpsC, velB, wetA, wsc1* |

Since the *Aspergillus* WetA proteins have a highly conserved putative DNA-binding domain and have conserved functions in the overall conidiation process, we hypothesized that *Afu*WetA and *Afl*WetA recognize a similar *Ani*WetA WRE. To test our hypothesis, we searched the *Ani*WetA WRE in the *A. fumigatus* and *A. flavus* genomes and summarized the results in Fig 6. Only 15 genes, including *wetA*, that contain a WRE in their upstream 1.5 kb regions in all three species also exhibit consistent differential expression (Table 5). We further searched for WRE occurrences in the 1.5 kb sequence upstream of other *wetA* orthologs in different *Aspergillus* and other fungal species and found that the WRE in the upstream region of *wetA* genes is completely conserved throughout the family Aspergillaceae (Fig 7).

**Table 5.** Conserved DEGs with a WRE in *Aspergilli*

|  | *A. nidulans ID* | *A. flavus ID* | *A. fumigatus ID* |
| --- | --- | --- | --- |
| **Up-regulated**<br>**in $\Delta wetA$ conidia** | AN1156 | AFLA_068310 | Afu1g11450 |
|  | AN10598 | AFLA_101220 | Afu3g07020 |
|  | AN6088 | AFLA_045760 | Afu2g09282 |
|  | AN4836 | AFLA_102180 | Afu3g07290 |
|  | AN3752 | AFLA_073850 | Afu7g04580 |
| **Down-regulated**<br>**in $\Delta wetA$ conidia** | AN4464 | AFLA_112150 | Afu4g07690 |
|  | AN1524 | AFLA_078640 | Afu8g05330 |
|  | AN5715 | AFLA_127800 | Afu1g06770 |
|  | AN10265 | AFLA_014960 | Afu2g15910 |
|  | AN10551 | AFLA_112710 | Afu4g07140 |
|  | AN5215 | AFLA_087840 | Afu6g07490 |
|  | AN8763 | AFLA_131750 | Afu6g02960 |
|  | AN9037 | AFLA_023560 | Afu3g15190 |
|  | AN4716 | AFLA_091920 | Afu5g08580 |
|  | AN1937 | AFLA_052030 | Afu4g13230 |

**Fig 6.**
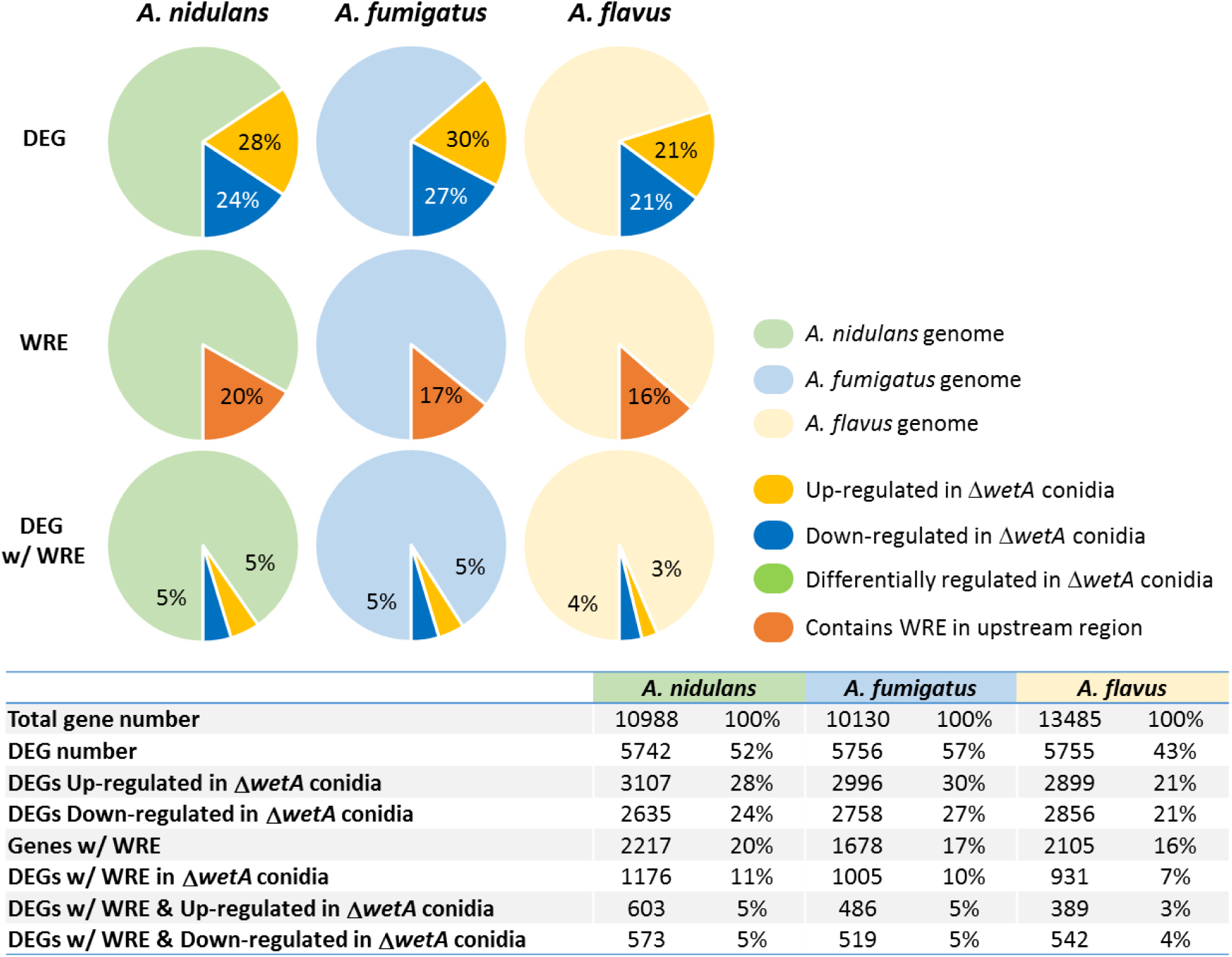
Overlap between DEGs and WRE-containing genes in three *Aspergillus* species. The percentages of genes differentially expressed in the Δ*wetA* conidia (DEG), the percentage of genes that contain predicted WRE sequences in their upstream 1.5 kb regions (WRE), and the DEGs with a WRE in their upstream 1.5 kb regions (DEG w/ WRE) are shown. The *A. nidulans, A. fumigatus*, and *A. flavus* genes are shown in light green, light blue, and light orange, respectively.

**Fig 7.**
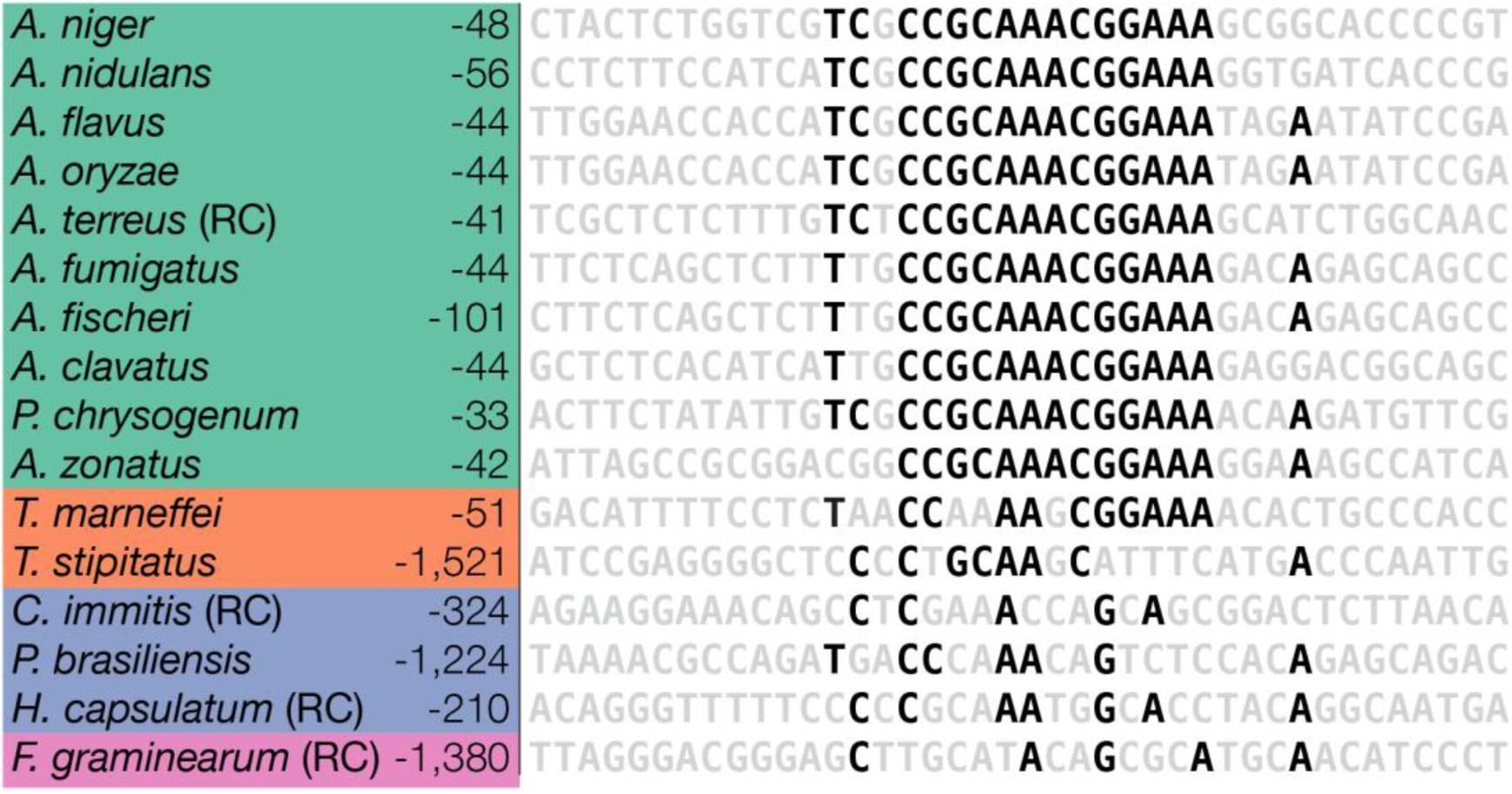
WRE occurrences in the upstream regions of *wetA* orthologs in representative fungi. WRE occurrences were identified in a series of regions located upstream of *wetA* orthologs. Numbers to the left of the sequence indicate at what position relative to the translation start site the sequence shown begins. The sequences shown are from 15 bp upstream of the WRE occurrence that was identified by FIMO (42) with the lowest p-value, to 14 bp downstream of the WRE occurrence. Bases are colored black if they are conserved in at least 60% of the species. Green – Aspergillaceae. Orange – Trichocomaceae. Blue – Onygenales. Purple – Sodariomycete. (RC) – Reverse Complement.

Taken together, we conclude that WetA regulates the *Aspergillus* conidial transcriptomes through both direct and indirect methods and controls species-specific GRNs to achieve conserved and diverged functions.

## Discussion

While WetA is well known as the key regulator of multiple cellular and chemical developmental processes in Ascomycetes (7, 8, 17–23, 30, 9–16), the regulatory mechanisms behind it employs are not known. In this study, we investigated the roles of WetA-mediated GRNs in the model organism *A. nidulans*, the human pathogen *A. fumigatus*, and the aflatoxin producer *A. flavus* and further identified a potential WetA binding motif in *A. nidulans*.

Previous studies suggested that *Ani*WetA is required for activating a set of genes whose products comprise, or direct the assembly of, the conidial wall layers and also ensure proper cytoplasmic metabolic remodeling including massive trehalose biogenesis (20, 22, 43). We also reported that *Afl*WetA is involved in the regulation of conidial secondary metabolism and hypothesized that this was done by WetA controlling a group of TFs and signaling pathways (30). Our RNA-seq results here show that 52%, 57%, and 43% of *A. nidulans, A. fumigatus,* and *A. flavus* transcriptomes were differentially regulated in the Δ*wetA* conidia respectively, suggesting a broad regulatory role for WetA in aspergilli (Table 2, Fig 6). While *Ani*WetA, *Afu*WetA, and *Afu*WetA are functionally conserved in many aspects of developmental processes in conidia, the specific genes regulated by WetA are divergent in each species. Although WetA regulates a large number of common orthogroups in aspergilli, only 9% of *Aspergillus* orthogroups were consistently all over-or underexpressed in Δ*wetA* conidia from the three species, suggesting that while the WetA-mediated regulation is functionally conserved, the WetA-mediated GRNs have been highly rewired (Fig 8).

**Fig 8.**
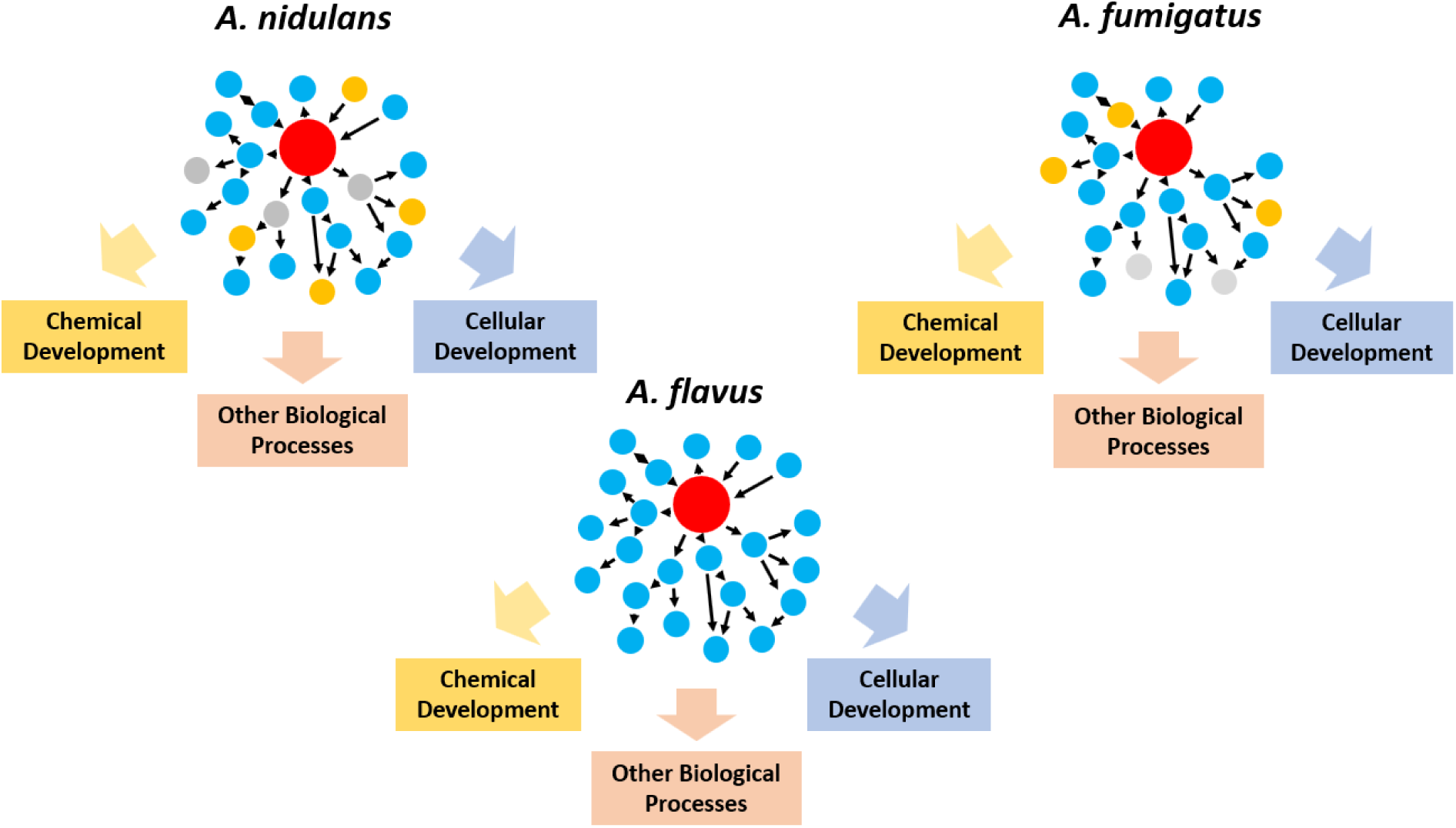
Proposed model of the rewired WetA-mediated GRNs in *Aspergilli*. We propose that WetA-mediated GRNs have been rewired during the evolution of *Aspergillus* species but that their regulatory logic (i.e., their regulation of chemical development, cellular development, and other biological processes) remains conserved. Red circle: WetA. Blue circle: Targets that are consistently up-/down-regulated by WetA. Yellow circle: Targets that are divergently regulated by WetA. Grey circle: Targets that are not regulated by WetA.

An example of a divergent WetA-mediated GRN whose output is the conserved regulation of a biological process is *Aspergillus* asexual development (Fig 3). In all three species, loss of *wetA* leads to increased levels of the central regulator *brlA* in conidia and shuts down asexual development. However, the set of regulatory events that result in WetA-mediated repression of *brlA* are unique to each species (Fig 3).

Although WetA shows broad regulatory effects in *Aspergillus* species, only 15 genes with the WRE in their upstream regions were consistently under-or overexpressed in the Δ*wetA* conidia. The list of peak-associated genes in *A. nidulans* includes *wetA* and the important developmental regulators *vosA* and *velB*, suggesting that these genes may play crucial roles in conidiation and thus be conserved during evolution. VosA and VelB are both members of the *velvet* family of proteins (44–46). Moreover, the VosA-VelB complex is a crucial functional unit controlling conidia maturation (45 –47). Loss of *vosA* causes some phenotypes similar to those by the loss of *wetA*, like a reduction in trehalose amount (48), suggesting that part of the WetA-mediated GRN may be controlled by regulating VosA. Previous studies show that *Ani*WetA contains an *Ani*VosA binding motif in its upstream 2 kb region (44), implicating the cross feedback regulation of WetA by VosA. Taken together, the WetA-mediated regulatory pathway may cross-talk with the *velvet* regulatory pathways via the cooperative activity of WetA/VosA/VelB.

We further examined the WetA-mediated GRNs controlling other pathways based on previously characterized, conserved WetA functions. First, we analyzed genes involved in conidial integrity for their WetA-regulation. The genes associated with trehalose biosynthesis are almost all underexpressed in all three of the Δ*wetA* conidia (Fig 4). Similarly, almost all the genes associated with β-(1,3)-glucan biosynthesis were overexpressed in all three of the Δ*wetA* conidia (Fig 4). These results explain the dramatically reduced amount of trehalose and increased content of β-(1,3)-glucan in the Δ*wetA* conidia (8, 30) and suggest a conserved WetA-mediated GRN for activation of trehalose biogenesis and repression of β-(1,3)-glucan biosynthesis. WetA’s function is likely diverged in α-(1,3)-glucan metabolism. *Ani*WetA upregulates the α-(1,3)-glucan synthase *Aniags2* but downregulates all the genes associated with α-(1,3)-glucan degradation except AN1604 (Fig 4). In contrast, *Afu*WetA downregulates all the α-(1,3)-glucan synthases (*Afuags1, Afuags2*, and *Afuags3*), but has mixed effects on the genes associated with α-(1,3)-glucan degradation in conidia (Fig 4). In conidia, *Afl*WetA shows mixed effects on both the genes associated with α-(1,3)-glucan biosynthesis and degradation (Fig 4).

WetA is involved in the regulation of hydrophobin expression. Only one of the five hydrophobin-encoding genes in *A. nidulans* was not differentially expressed in the Δ*wetA* conidia, and only *AnidewA* was underexpressed (Fig 5). In *A. fumigatus*, all six hydrophobin-encoding genes were overexpressed in the Δ*wetA* conidia (Fig 5). In *A. flavus*, three of five hydrophobin-encoding genes were underexpressed in the Δ*wetA* conidia, one of them was not regulated, and only *AflrodA* was up-regulated (Fig 5). Since the loss of *wetA* causes lower hydrophobicity of conidia, there might be other unidentified hydrophobins controlled by *Afu*WetA.

*Afu*WetA is diverged relative to *Ani*WetA and *Afl*WetA in its regulation of melanin biosynthesis. A previous study showed that *wA*, the first regulator in the DHN-melanin synthesis pathway, is activated by WetA in *A. nidulans* conidia (20, 49). Our RNA-seq analyses have revealed that both *AniwA* and *AflwA* were underexpressed in the Δ*AniwetA* and Δ*AflwetA* conidia (Fig 5).Moreover, *Aflayg1*, the second gene in the DHN-melanin pathway (50) was underexpressed in the Δ*AflwetA* conidia (Fig 5). Surprisingly, although the Δ*AfuwetA* conidia are colorless, all theDEGs associated with both DHN-melanin and pyomelanin biosynthesis were overexpressed in the Δ*AflwetA* conidia (Fig 5), suggesting the melanin biosynthesis pathway in *A. fumigatus* may have uniquely evolved.

We identified a potential WRE (5’-CCGYTTGCGGC-3’), which is highly similar to the *Saccharomyces cerevisiae* Ixr1, Dal81, and Leu3 motifs (51–53). Although 53% of genes in the *A. nidulans* genome were differentially regulated in the Δ*AniwetA* conidia, only 21% of them contain a WRE in their upstream 1.5 kb regions (Fig 6), suggesting that *Ani*WetA might serve as a conserved regulatory hub which controls a group of regulators of various biological processes. Our data support a model where the WetA-mediated regulation is carried out via both direct and indirect interactions to control a downstream cascade of genes (Figure 9). We also scanned the *A. fumigatus* and *A. flavus* genomes for instances of the WRE and found that, while similar numbers of genes contained the WRE compared to *A. nidulans*, the makeup of that list of genes was different.

In conclusion, our studies provide the first clear and systematic dissection of WetA, an evolutionarily and functionally conserved regulator of morphological and chemical development of filamentous fungal conidiation. Moreover, we have revealed the molecular mechanisms of WetA as a DNA-binding, multi-functional regulator governing the diverged processes of cellular differentiation, chemical development, and cell survival across a genus of filamentous fungi, which advances our knowledge of spore formation in pathogenic and toxigenic fungi.

## Materials and methods

### Strains, media, and culture conditions

All strains used in this study are listed in Table S1. The fungal strains were grown on minimal medium (MM) with appropriate supplements as described (48, 54), and incubated at 37°C (*A. nidulans* and *A. fumigatus*), or 30°C (*A. flavus*). For liquid cultures, conidia were inoculated in liquid MM and incubated at 37°C or 30°C, 220 rpm. Conidiation induction was performed as described (55).

### Generation of *wetA* deletion and complemented strains

We generated the deletion (Δ) and complement (C’) strains of *wetA* in *A. nidulans* (*AniwetA*). The oligonucleotides used in this study are listed in Table S1. Briefly, the deletion construct containing *A. fumigatus pyrG* marker with 5’ and 3’ flanking regions of *AniwetA* were introduced into the recipient strain RJMP1.59 (56). To generate complemented strains, a WT *AniwetA* gene region, including its 2 kb upstream region, was cloned to pHS13 (45). The resulting plasmid pMY1 was then introduced into the recipient Δ*AniwetA* strain TMY3, resulting in isolation of TMY4. Multiple Δ*AniwetA* strains were generated and all behaved the same in every assay. Multiple C’ *AniwetA* strains were generated and all behaved identically to one another as well. The Δ*AfuwetA* (TSGw4), Δ*AflwetA* (TMY1), and C’*AflwetA* (TMY2) strains were generated in previous studies (8, 30).

### Nucleic acid manipulation

The genomic DNA and total RNA isolation for Northern blot analyses was performed as described (55, 57, 58). For RNA-seq and ChIP-seq, fresh conidia from 2-day-old solid cultures grown at 37°C or 30°C of WT and Δ*wetA* strains were harvested.

### RNA sequencing

Total RNA from 4 *A. nidulans* biological replicates, 3 *A. flavus* biological replicates, and 3 *A. fumigatus* biological replicates was extracted and submitted to ProteinCT Biotechnologies (Madison, WI) and the University of Wisconsin Gene Expression Center (Madison, WI) for library preparation and sequencing. For each replicate, a strand-specific library was prepared from total RNA using the Illumina TruSeq Strand-specific RNA sample preparation system. All replicates’ libraries were sequenced (PE100bp for *A. nidulans* and SE100bp for *A. fumigatus* and *A. flavus*) using the Illumina HiSeq2500.

The *A. flavus* expression data were analyzed as previously reported (30). The following analyses were carried out for the *A. fumigatus* and *A. nidulans* data. The overall quality of the raw sequence reads was verified using version 0.11.5 of FastQC (59). The genomes and annotation were downloaded from FungiDB and used for mapping (60). Mapping of the raw sequence reads to the genome was carried out with version 2.1.1 of Tophat2 (61), and the default settings were used except that the max intron length was set to 4,000 bases. The alignment files were compared against the gene annotation file, and raw counts for the number of reads mapping to each gene were generated using version 0.6.1p1 of HTSeq-count (62). Differential expression analysis of the raw counts was carried out using version 1.14.1 of DESeq2 (63). Genes were considered differentially expressed between the WT and Δ*wetA* conidia if their adjusted p-value was less than 0.05 and their log2 fold-change was smaller than-1 or greater than 1. All RNA-seq data files are available from the NCBI Gene Expression Omnibus database (*A. nidulans* and *A. fumigatus*: GSE114143; *A. flavus*: GSE95711).

### Functional Enrichment and Orthogroup identification

Gene Ontology enrichment analyses were carried out using the tool available at FungiDB (60). Unless otherwise stated, default settings were used in FungiDB, and redundant terms were collapsed with the REVIGO tool (64) using the “Tiny” setting for allowed similarity. Orthologs were identified using OrthoMCL with the settings: p-value cutoff of 6e-6, percent identity cut-off of 30%, percent match cut-off of 70%, MCL inflation value of 2, and Maximum weight allowed of 180.

### Chromatin immunoprecipitation sequencing (ChIP-seq)

ChIP assays were performed using MAGnify ChIP assays (Invitrogen) according to the manufacturer’s instructions. Briefly, 10^9^ of *A. nidulans* WT conidia were cross-linked with 1% formaldehyde, lysed and broken as described (65). Cell lysates were sonicated to shear DNA to 300-500 bp and were immunoprecipitated with the rabbit anti-WetA polyclonal antibodies (GenScript, NJ). Two experiments were performed, each with biological triplicates. In the first experiment, 10% of the supernatants was kept as an input control (input represents PCR amplification of the total sample) and compared to the ChIP sample. In the second experiment, the ChIP sample from the WT strain was compared to the ChIP sample from the Δ*wetA* strain. ChIP DNA samples were sent for ChIP-Seq service (ProteinCT, WI). Libraries were prepared using the TruSeq ChIP Library Preparation Kit (Illumina, CA) and sequenced on a HiSeq2500 with single reads of 50 bp. Approximately 8-30 M reads were achieved per replicate.

ChIP-seq reads were first trimmed using version 0.36 of the Trimmomatic software (66) and then version 0.7.15 of the BWA-MEM software (67) was used to map reads to the *A. nidulans* (FGSC A4) genome. Reads with any of the following flags were removed: unmapped, secondary alignment, or supplementary mapped read. Reads with a mapping quality (MAPQ) score of 0 were also removed. Duplicate reads were removed and samples were pooled using version 1.3 of the SAMtools software (68). Version 2.1.1.20160309 of the MACS2 software (69) with the settings -g 2.93e7 –s 101 --nomodel --extsize was used to call peaks. Extension sizes were calculated using SPP (70, 71). Peaks that exhibited a fold-change greater than 2 and a q-value less than 0.001 were used in further analyses. Peak lists were combined from each of the ChIP experiments. The ChIP-seq data is available from the NCBI Gene Expression Omnibus database (GSE114141).

### Motif discovery analyses

To discover the WetA-Response Element (WRE), 100 bp of sequence surrounding the summits of the 157 combined peaks were pulled from the *A. nidulans* genome using the bedtools software, version 2.26.0 (72) and submitted to the MEME-ChIP software, version 4.12.0 (41). MEME was instructed to search for 10 motifs, 5-21 bp in length; all other settings were left at default. Instances of the WRE were identified in the upstream regions (1.5 kb upstream of the translation start) of all genes in the three *Aspergillus* genomes using the FIMO software (42) with a p-value cutoff of 5e-5.

